# Structural basis of ribosomal RNA repair by RtcB2

**DOI:** 10.64898/2026.11.03.763826

**Authors:** Yannan Tian, Qixin Zhou, Qianxi Yu, Yini Huang, Wenju Zhang, Jun Huang, Ting Yu, Fuxing Zeng

## Abstract

Ribotoxins disable ribosomes by cleaving ribosomal RNA, but how repair enzymes recognize and restore these damaged machines remains unclear. Here, we report three cryo–electron microscopy structures of the RNA ligase RtcB2 bound to damaged *Escherichia coli* 30S subunits. An extended interface stabilizes the ribosomal head and decoding-center scaffold while positioning the cleaved 3′-phosphate end within the catalytic pocket. Two structures contain enzyme-linked guanosine monophosphate, whereas the third captures its transfer to the RNA. RNA activation accompanies RtcB2 reorientation and entry of the 5′-hydroxyl-containing segment into an adjacent cleft. Comparisons with assembled ribosomes reveal structural barriers to substrate access. These structures connect recognition of a damaged ribosome to chemical activation of its RNA for repair.

## INTRODUCTION

Ribosomes are targets of ribotoxins that cleave rRNA and disrupt protein synthesis (*1–3*). Although such damage can inactivate the ribosome, cleavage does not necessarily commit the particle to degradation. The RNA ligase RtcB can reconnect the MazF-truncated 3 ′ terminus of 16S rRNA (*4*), and the specialized RtcB2–PrfH system rescues and repairs ribosomes cleaved within the decoding center (*5*). These findings establish RNA repair as a route to recovering damaged ribosomes. However, how a repair ligase recognizes a lesion within the extensive RNA–protein architecture of a ribosomal subunit and positions the broken ends for ligation remains unresolved.

RNA ligases restore phosphodiester bonds between broken RNA ends. Classical RNA ligases join 3 ′ - hydroxyl and 5′-phosphate termini (*6–8*), whereas RtcB, identified as an RNA ligase in bacteria, archaea, and mammals in 2011, joins 3′-phosphate and 5′-hydroxyl termini (*9–11*). RtcB reacts with GTP to form a covalent enzyme–GMP intermediate through an active-site histidine (*12–15*). Substrates bearing a 2′,3′-cyclic phosphate undergo conversion to a 3 ′ -phosphate before guanylate transfer generates an RNA(3′)pp(5′)G intermediate. Subsequent attack by the 5′-hydroxyl forms a 3′–5′ phosphodiester bond and releases GMP (*14, 16*). Structural studies have defined the catalytic architecture and nucleotide-bound states of RtcB (*12, 13, 15*), the organization of the metazoan tRNA ligase complex (*17, 18*), and its regulation by Archease and PYROXD1 (*19, 20*). A structure of RtcB bound to a six-nucleotide DNA oligonucleotide has further revealed contacts with a 5′-hydroxyl terminus (*21*). Nevertheless, how RtcB engages a damaged ribosome and couples recognition of this macromolecular substrate to activation of the cleaved RNA remains unknown.

Bacterial RtcB paralogs can exhibit distinct substrate preferences. In *Escherichia coli*, RtcB1 repairs cleaved tRNA, whereas RtcB2 specifically repairs the decoding-center lesion between A1493 and G1494 of 16S rRNA (*5*). We previously identified the *rtcB2*–*prfH* operon as a specialized ribosome rescue and repair system present in a subset of bacteria (*5*). PrfH, a homolog of the class I release factors RF1 and RF2, recognizes damaged ribosomes and catalyzes peptidyl-tRNA hydrolysis (*5, 22*). These findings led to a model in which peptide release and subsequent subunit dissociation enable RtcB2 to access the damaged 30S subunit (*5*). More recently, RtcB2-mediated repair was reported in preparations of damaged 70S complexes containing mRNA and P-site tRNA (*23*), raising questions about the ribosomal states that permit repair. Defining the RtcB2–ribosome interface is therefore essential to understanding both substrate selectivity and access to the damaged decoding center.

Here, we report three cryo–electron microscopy (cryo-EM) structures of *E. coli* RtcB2 bound to ribotoxin-cleaved 30S subunits. The structures reveal an extended recognition interface that positions the damaged rRNA within the catalytic center. Two states contain GMP covalently linked to the catalytic histidine, whereas the third contains the guanylylated rRNA intermediate, capturing states before and after guanylate transfer to the cleaved 3′-phosphate end. RNA activation is accompanied by reorientation of RtcB2 and engagement of the 5 ′ -hydroxyl-containing segment in an adjacent cleft. Comparisons with unbound damaged subunits and assembled 70S ribosomes further support conformational stabilization associated with RtcB2 binding and define structural constraints on substrate access. Together, these findings provide, to our knowledge, the first structures of an RNA repair ligase engaged with a damaged ribosomal subunit and reveal how ribosomal substrate recognition is coupled to RNA-end activation.

## RESULTS

### Overall Structures of *Ec*RtcB2 Bound to Damaged 30S Subunits

To investigate how RtcB2 engages its ribosomal substrate, we reconstituted a complex of purified *E. coli* RtcB2 (*Ec*RtcB2) with damaged 30S subunits (d30S) isolated from cells expressing the ribotoxin CdiA-CT^ECL^, which cleaves 16S rRNA between A1493 and G1494 within the decoding center (*2, 5*). Single-particle cryo-EM analysis of three combined datasets, followed by focused 3D classification of the *Ec*RtcB2-binding region, resolved three *Ec*RtcB2–d30S states, designated *Ec*RtcB2–d30S-I, *Ec*RtcB2–d30S-II, and *Ec*RtcB2–d30S-III, at overall resolutions of 2.68, 2.65, and 2.76 Å, respectively (F**ig. 1** **and Figs. S1 and S2**).

**Fig. 1.**
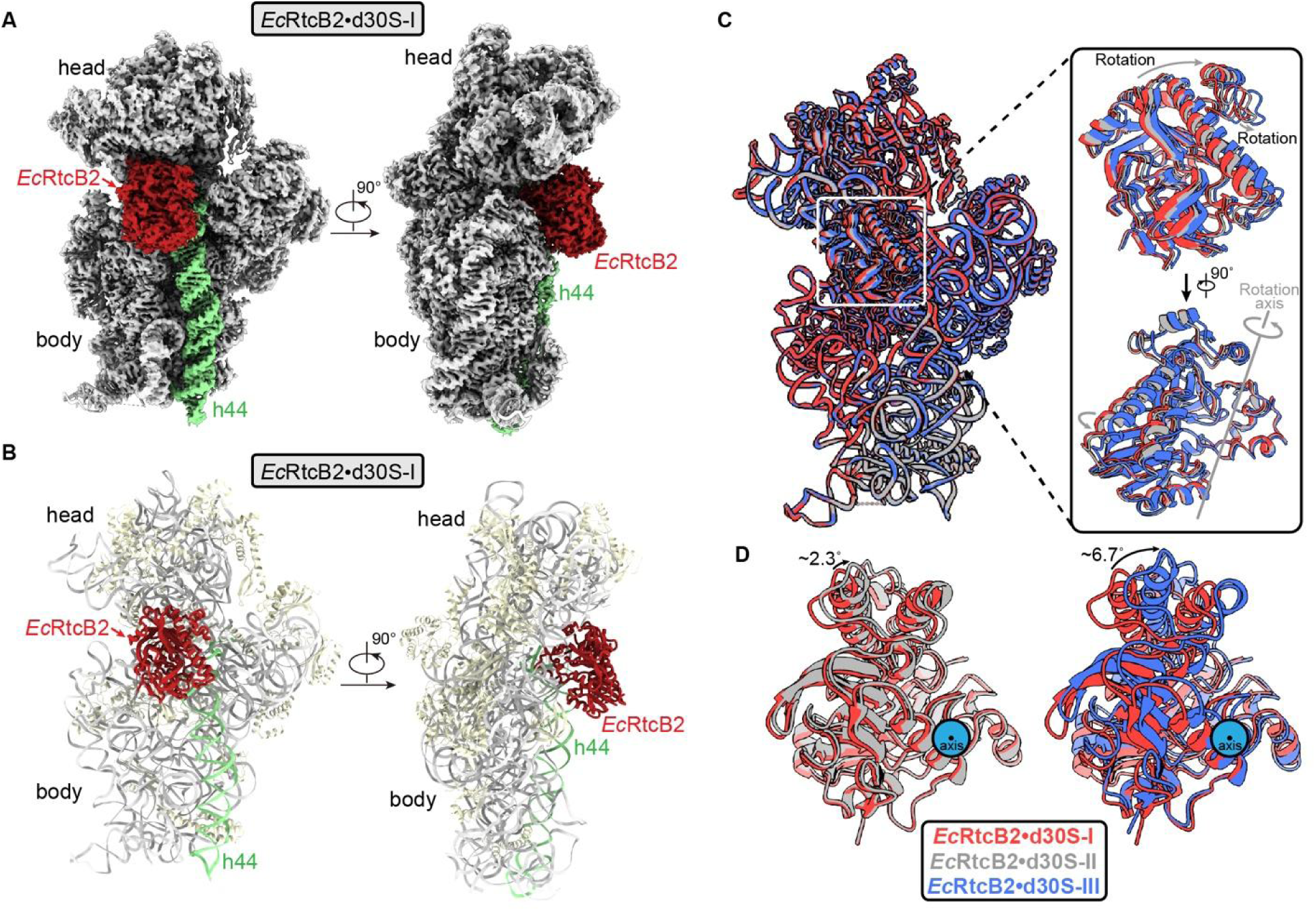
Overall structures of *Ec*RtcB2 bound to damaged 30S subunits. (**A** and **B**) Cryo-EM map (**A**) and atomic model (**B**) of *Ec*RtcB2•d30S-I, shown in two views related by a 90° rotation. *Ec*RtcB2 is colored red and h44 green. The 30S head and body are labeled. (**C**) Superposition of states I (red), II (gray), and III (blue) after alignment on the 30S body. Enlarged views show differences in *Ec*RtcB2 orientation, with the approximate rotation axis indicated. (**D**) Pairwise comparisons of *Ec*RtcB2 in states I and II (left) and states I and III (right), retaining the body-aligned coordinates. The corresponding rotations are approximately 2.3° and 6.7°, respectively.

All three structures show *Ec*RtcB2 bound to the intersubunit face of the 30S subunit, at the junction between the head and body (**Fig. 1A, B**). The enzyme engages a composite surface formed by 16S rRNA and ribosomal protein uS12, positioning its catalytic pocket adjacent to the cleaved region of h44. Secondary-structure elements are numbered according to the experimentally determined *Ec*RtcB2 structure and are annotated above a sequence alignment with *Ec*RtcB1, *Ec*RtcB3, and *Pho*RtcB (**Fig. S3**).

Alignment on the 30S body reveals nearly identical head orientations across the three bound states (**Fig. 1C**). *Ec*RtcB2 retains the same overall binding location but differs in orientation, with rotations of approximately 2.3° and 6.7° between states I and II and between states I and III, respectively (**Fig. 1D**). These structures therefore reveal limited reorientation of *Ec*RtcB2 within a largely preserved ribosomal framework. The same datasets also yielded five unbound d30S reconstructions, designated states A–E, providing structural references for examining the head and h44 in the absence of bound *Ec*RtcB2 (**Figs. S1 and S4**).

### Recognition of the Damaged 30S Subunit by *Ec*RtcB2

*Ec*RtcB2 engages the damaged 30S subunit through an extended interface encompassing the decoding center, head-associated elements of 16S rRNA, and ribosomal protein uS12 (**Fig. 2**). Predominantly electropositive surfaces coincide with the rRNA-contacting regions, including the pocket accommodating A1492, whereas the uS12 interface combines nonpolar packing with polar contacts (**Fig. S5**). This composite interface connects recognition of the ribosomal scaffold with positioning of the cleaved rRNA.

**Fig. 2.**
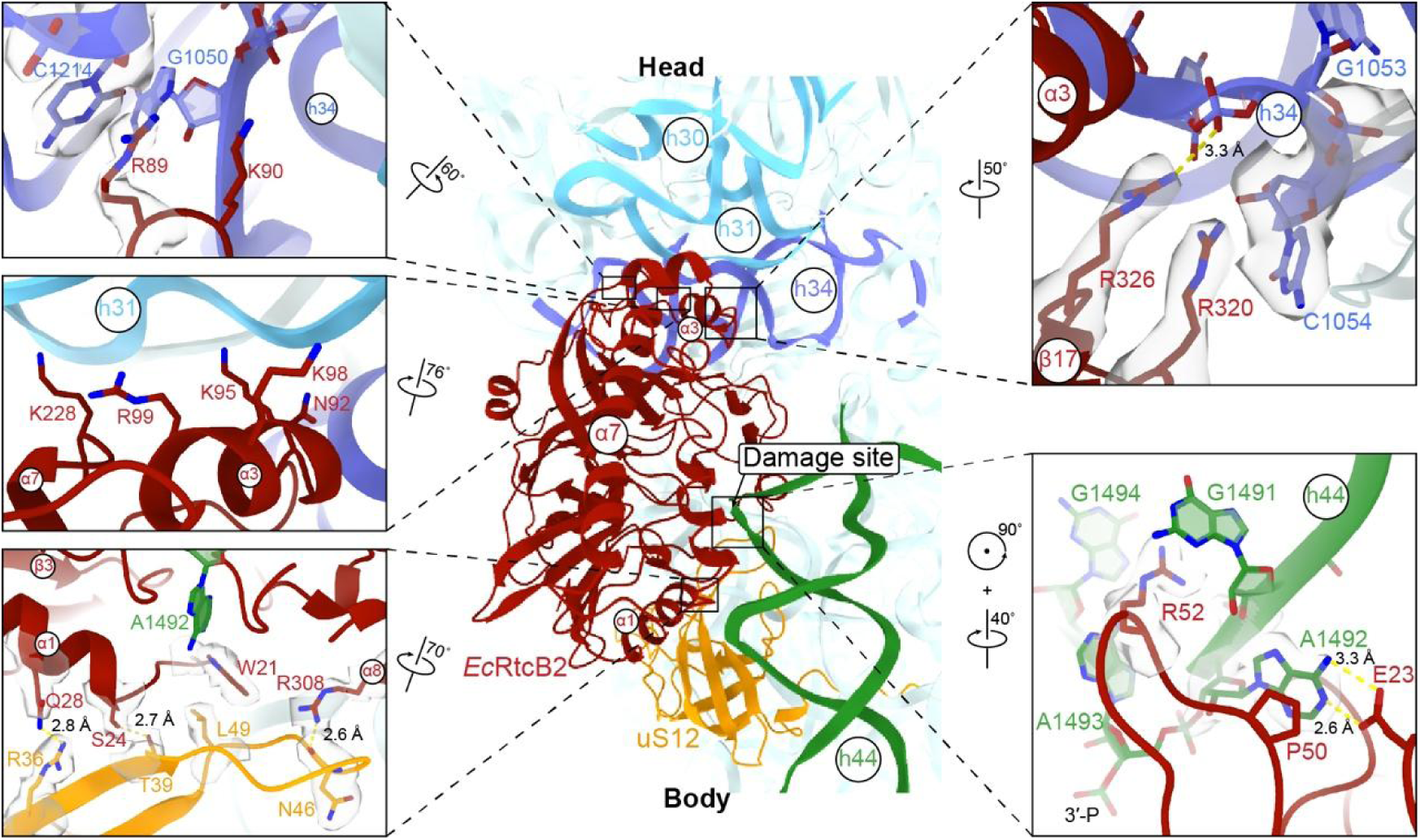
Recognition of the damaged 30S subunit by *Ec*RtcB2. The central view shows *Ec*RtcB2 (red) contacting head-associated 16S rRNA helices, h44 (green), and uS12 (orange). Enlarged views show key contacts at the enzyme–ribosome interface, with selected residues and nucleotides displayed as sticks and corresponding cryo-EM map features as semitransparent surfaces. Dashed yellow lines indicate interatomic contacts, with distances labeled. Viewing rotations are indicated.

At the damaged decoding center, *Ec*RtcB2 contacts the rRNA segment immediately upstream of the cleavage between A1493 and G1494. R52 inserts into h44, with its guanidinium group packing against the G1491 base, while A1492 projects into a complementary pocket on *Ec*RtcB2, where it packs against P50 and forms hydrogen-bonding contacts with E23 (**Fig. 2**). This interlocking interface may stabilize the upstream rRNA segment, with the adjacent A1493 nucleotide accommodated within the catalytic pocket and its terminal 3′-phosphate positioned at the active site. By contrast, the G1494-containing 5′-hydroxyl end remains outside the catalytic center. The observed structure therefore captures engagement of the 3′-phosphate end without juxtaposition of both reactive termini in a configuration suitable for end joining.

On the 30S head, R320 packs against the C1054 base of h34, a contact retained in all three bound states, while the neighboring R326 contacts the phosphate backbone of G1053 (**Fig. 2**). Additional contacts involve K90 and the G1050 phosphate, together with packing of R89 against the C1214 base. A neighboring surface containing K90, N92, K95, R99, and K228 engages h31. Toward the 30S body, H49, H137, R172, K257, and K306 form an additional contact surface along h44 near C1397 and C1400, with K306 also contacting h28 (**Fig. S5**). These contacts distribute recognition across several head-associated rRNA elements rather than confining binding to the cleavage site.

*Ec*RtcB2 also contacts uS12 through an interface involving W21, S24, Q28, and R308, with uS12 residues T39, R36, N46, and L49 contributing to the opposing surface (**Fig. 2**). Packing involving W21 and uS12 L49 is accompanied by a network of polar contacts across the interface. Together, the contacts with uS12 and head-associated rRNA position *Ec*RtcB2 across the head–body junction, while the interlocking interface around G1491 and A1492 supports local accommodation of the cleaved RNA. This arrangement suggests that productive substrate engagement depends on both the damaged RNA terminus and its presentation within the ribosomal scaffold.

### Conformational Variation in Unbound and *Ec*RtcB2-Bound d30S

To examine conformational differences associated with *Ec*RtcB2 engagement, we compared the five unbound d30S reconstructions with the bound structures after alignment on the 30S body, using *Ec*RtcB2•d30S-I as the reference. The unbound states displayed distinct head orientations, with rotations ranging from 3.83° to 19.30° and shifts of 3.72–13.17 Å relative to the bound reference (**Table 1**). Variation was also apparent near the h44 cleavage site, where displacement of the selected reference position ranged from 6.22 Å to approximately 18.38 Å. Several of these distances could only be estimated because the corresponding map features were poorly resolved (**Table 1**). These comparisons reveal conformational heterogeneity in the head and the damaged region of h44 in the absence of bound *Ec*RtcB2.

**Table 1.** Structural Differences in the 30S Head and G1494 Phosphate Relative to *Ec*RtcB2–d30S-I.

|  |  | 30S head |  | G1494 phosphate |
|---|---|---|---|---|
| Reconstructions | States | Rotation (˚) | Shift (Å) | Shift (Å) |
| *Ec*RtcB2•d30S | I | - | - | - |
|  | II | 0.60 | 0.13 | 0.02 |
|  | III | 0.90 | 0.30 | 5.67 |
| unbound d30S | A | 12.64 | 8.01 | 6.22 |
|  | B | 9.40 | 4.91 | ~18.38<sup>a</sup> |
|  | C | 9.10 | 6.30 | ~9.95<sup>a</sup> |
|  | D | 3.83 | 3.72 | ~8.97<sup>a</sup> |
|  | E | 19.30 | 13.17 | ~15.63<sup>a</sup> |
| unbound 70S |  | 15.60 | 11.91 | 6.73 |

<sup>a</sup>Because the cleavage-site region is poorly resolved in these maps, displacements were estimated from the corresponding map features.

By contrast, the three *Ec*RtcB2-bound structures shared nearly identical head configurations. Relative to state I, the heads in states II and III differed by only 0.60° and 0.90°, with shifts of 0.13 and 0.30 Å, respectively (**Fig. 1C and Table 1**). *Ec*RtcB2 itself underwent larger reorientations of approximately 2.3° and 6.7°, respectively (**Fig. 1D**). Thus, differences in enzyme orientation were accommodated without appreciable movement of the ribosomal head. The h44 scaffold was also broadly preserved, although local rearrangement occurred at the cleavage site in state III, as described below.

Together, these comparisons support stabilization of the 30S head and the overall h44 framework upon *Ec*RtcB2 binding. The extended interface involving head-associated rRNA, h44, and uS12 provides a structural basis for constraining the ribosomal scaffold while permitting local changes associated with RNA-end activation.

### Structural Snapshots Before and After Guanylate Transfer to the Cleaved rRNA

The three *Ec*RtcB2•d30S structures reveal distinct covalent states of the ligation pathway. In states I and II, GMP is covalently linked to the catalytic residue H294, whereas in state III it is attached to the terminal 3′-phosphate of A1493 (**Fig. 3A,B and Fig. S6**). These structures capture the enzyme–GMP and RNA–GMP intermediates before and after guanylate transfer to the cleaved rRNA (*12, 14*).

**Fig. 3.**
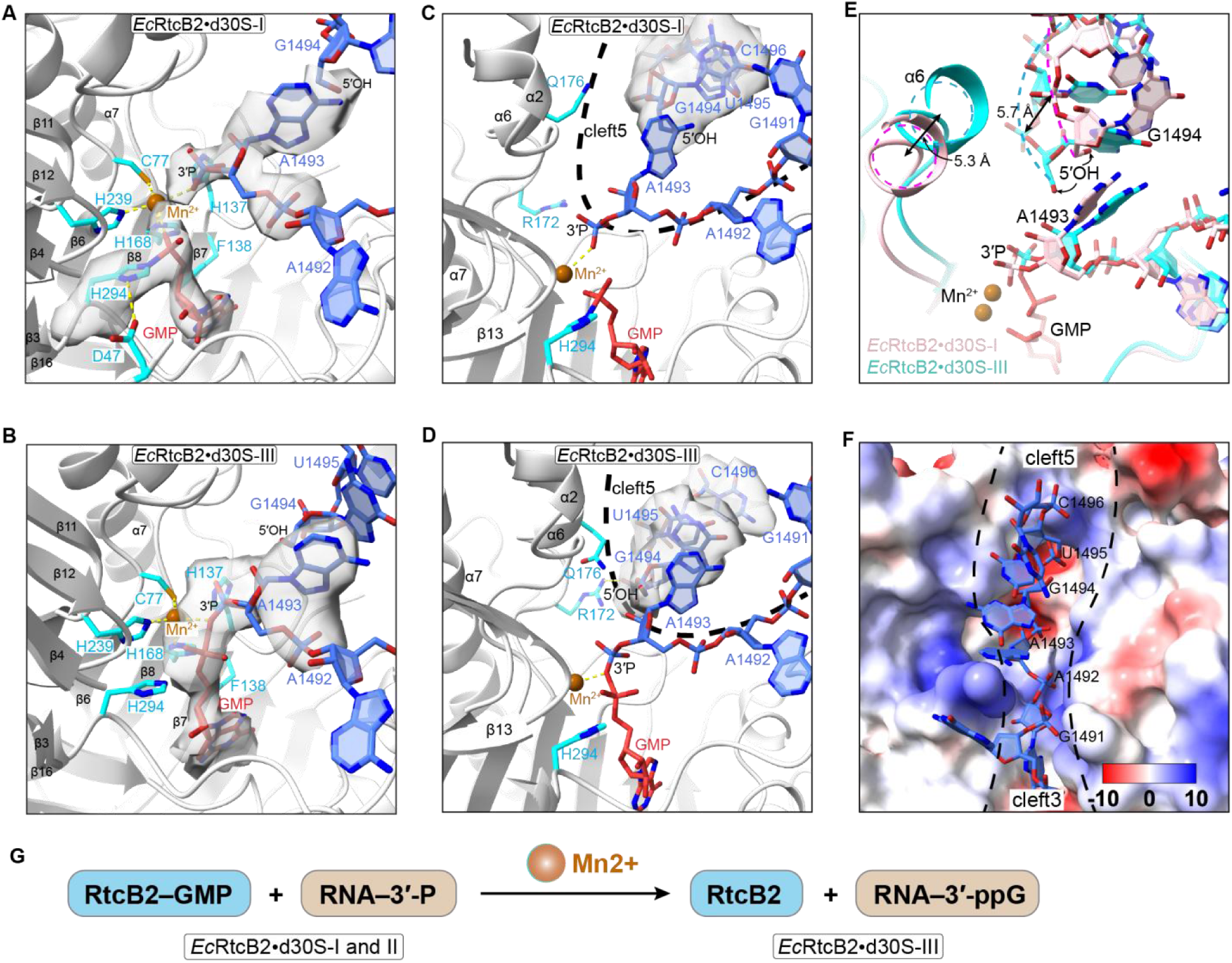
Structural snapshots before and after guanylate transfer to the cleaved rRNA. (**A** and **B**) Active-site configurations in states I (**A**) and III (**B**), showing GMP linked to H294 or the A1493 3 ′ -phosphate, respectively. Selected residues and nucleotides are shown as sticks with corresponding cryo-EM map features displayed as semitransparent surfaces. Mn^2+^ is shown as an orange sphere. (**C** and **D**) Accommodation of the cleaved rRNA in states I (**C**) and III (**D**). (**E**) Superposition of states I (pink) and III (cyan), highlighting local displacements. (**F**) Electrostatic surface of *Ec*RtcB2 in state III, showing the rRNA-binding clefts. Potentials range from −10 (red) to +10 (blue) kcal/(mol·e). (**G**) Reaction scheme assigning states I/II and III to the enzyme–GMP and RNA–GMP intermediates, respectively.

States I and II share the same chemical configuration despite their small difference in enzyme orientation. The active-site maps support a phosphoramidate linkage between H294 Nε2 and the GMP phosphorus (**Fig. 3A and Fig. S6**). H294 Nδ1 lies 3.44 Å in state I and 3.35 Å in state II from a side-chain carboxyl oxygen of D47, consistent with a hydrogen bond. This arrangement preserves the Asp–His interaction observed in PhoRtcB, where the corresponding residues D65 and H404 participate in histidine guanylylation (*12, 15*).

The A1493-containing 3′-phosphate end occupies a substrate-binding cleft, designated cleft3, adjacent to the H294-linked GMP. The metal ion, modeled as Mn^2+^, adopts an observed five-coordinate geometry involving the sulfur atom of C77, the Nε2 atoms of H168 and H239, and phosphate oxygens from GMP and the terminal 3′-phosphate of A1493 (**Fig. 3A and Fig. S6**). This coordination environment brings the enzyme-linked GMP and its RNA phosphate acceptor into proximity. The corresponding protein ligands are conserved in *Pho*RtcB, where C98, H234, and H329 contribute to the metal-binding center (*12, 15*). Whereas sulfate ions in earlier structures were used to infer substrate phosphate positions (*12*), the *Ec*RtcB2 complex directly locates a cleaved rRNA 3′-phosphate alongside the guanylylated enzyme.

State III exhibits a different covalent connectivity. The active-site map supports an RNA(3′)pp(5′)G linkage between the A1493 terminal phosphate and GMP, replacing the H294–GMP linkage observed in states I and II (**Fig. 3B**). The distance between H294 Nε2 and the GMP phosphorus increases to 3.35 Å, while the activated RNA terminus is accommodated within a catalytic pocket whose overall architecture remains largely conserved. The metal-coordination environment remains similar to that in states I and II, indicating that the change in covalent connectivity is accommodated within a largely preserved metal-binding architecture. These features identify state III as the product of guanylate transfer to the rRNA 3′-phosphate.

The change in chemical state is accompanied by increased engagement of the opposite side of the cleavage site. In state III, nucleotides G1494–C1496 extend into a second cleft, designated cleft5, that is not occupied by this segment in states I and II (**Fig. 3C,D**). R172 and Q176 contact the RNA backbone near G1494–U1495, while patches of positive electrostatic potential provide a complementary environment for the incoming segment (**Fig. 3D,F**). Comparison of states I and III reveals local displacements around helix α6 and the G1494-containing segment (**Fig. 3E**), accompanying reorientation of *Ec*RtcB2 within the largely unchanged ribosomal head configuration.

Together, the structures connect transfer of GMP from H294 to A1493 with a distinct binding configuration of the downstream rRNA. State III therefore represents a post-guanylate-transfer intermediate with increased engagement of the 5 ′ -hydroxyl-containing rRNA segment, preceding the final strand-joining reaction (**Fig. 3G**).

### Subunit Separation Permits RtcB2 Access to the Damaged Decoding Center

Our previous studies supported a sequential rescue-and-repair pathway in which PrfH-mediated peptidyl-tRNA hydrolysis precedes ribosome recycling and RtcB2-mediated repair of the damaged 30S subunit (*5, 22*). More recently, RtcB2-dependent repair was reported in reactions containing damaged 70S ribosomes, including mRNA-bound complexes carrying P-site tRNA (*23*). We therefore examined whether the *Ec*RtcB2-binding mode observed on d30S is compatible with an assembled 70S ribosome.

To assess whether the observed RtcB2-binding mode is compatible with an associated 70S ribosome, we aligned the 30S body of our RtcB2•d30S complex with that of a classical, nonrotated *E. coli* 70S complex containing mRNA and A- and P-site tRNAs (PDB: 7K00) (*24*). This comparison revealed steric overlaps with both tRNAs and the mRNA (**Fig. S7A**). Independently of these tRNA clashes, the α6 of RtcB2 overlapped helix H69 of the 23S rRNA, which projects into the intersubunit interface (**Fig. S7A**). Thus, both tRNA occupancy and the large subunit obstruct the RtcB2-binding configuration observed in the d30S complex.

We next examined the d70S reconstruction obtained from the same cryo-EM datasets (**Figs. S1 and S4F**). A discontinuity in the 16S rRNA backbone between A1493 and G1494 was consistent with cleavage at the targeted site, whereas no discernible map features attributable to RtcB2 were resolved. Placement of *Ec*RtcB2 by 30S-body alignment retained its overlap with H69 and revealed an additional overlap with the 30S head (**Fig. S7B**). Relative to *Ec*RtcB2–d30S-I, the d70S head differed by a rotation of 15.60° and a shift of 11.91 Å (**Table 1**), adopting an orientation that further restricted the space available for ligase binding (**Fig. S7B**). The observed RtcB2-binding mode therefore requires clearance of the intersubunit interface and a compatible 30S head conformation.

Together, these comparisons show that the observed *Ec*RtcB2-binding mode is incompatible with the examined 70S configurations. Subunit separation provides a route to remove the large-subunit barrier and permit engagement of a damaged 30S subunit with a compatible head conformation.

### Structural Features Associated with Substrate Selectivity in RtcB Ligases

Previous biochemical studies established distinct substrate preferences for *Ec*RtcB1 and *Ec*RtcB2, with the former repairing cleaved tRNA and the latter repairing damaged 30S subunits (*5*). To examine the structural basis of this distinction, we compared the *Ec*RtcB2•d30S complex with a tRNA model and with other RtcB structures.

Superposition of a helical segment of tRNA onto h44 placed corresponding tRNA bases near the *Ec*RtcB2 residues P50 and R52, in positions compatible with analogous packing interactions. The modeled tRNA could also be accommodated without apparent steric clashes with *Ec*RtcB2 (**Fig. S8**). Thus, this superposition did not reveal a steric barrier that would, by itself, explain the inability of *Ec*RtcB2 to repair tRNA. Instead, it indicated that local accommodation of an RNA helix and contacts near the cleavage site are insufficient to account for the observed substrate preferences.

We therefore examined the more extensive ribosomal recognition surface. *Ec*RtcB2 contacts both head-associated rRNA elements and uS12, extending its interface beyond the RNA segment accommodated near the catalytic pocket. Notably, the contact between R320 and C1054 in the 30S head is retained in all three *Ec*RtcB2-bound states (**Fig. 2**). Comparison with *Ec*RtcB1 revealed differences at the corresponding ribosome-facing surfaces, including substitutions at positions contributing to the head-associated rRNA and uS12 interfaces (**Fig. S3**). These differences suggest that conservation of the catalytic center does not imply conservation of the surface required to engage the damaged 30S subunit.

Structural comparison further revealed a similar overall catalytic-pocket architecture in *Ec*RtcB2, *Ec*RtcB1, and *Pho*RtcB, together with a difference in an adjacent loop (**Fig. S9**). *Ec*RtcB1 and *Pho*RtcB contain a longer loop near the region occupied by the 5′-hydroxyl-containing DNA strand in the *Pho*RtcB complex (PDB: 7LFQ) (*21*), whereas the corresponding loop in *Ec*RtcB2 is shorter. This difference identifies a potential contribution to substrate accommodation on the 5′-hydroxyl side of the active site. Together, these comparisons suggest that substrate selectivity reflects differences distributed across the RNA-binding surface, encompassing both peripheral substrate contacts and regions surrounding the reactive termini. The longer loop provides a candidate determinant of tRNA recognition, although its functional contribution remains unresolved.

## DISCUSSION

Repair allows damaged ribosomes to be recovered without replacing the entire ribonucleoprotein particle. Re-ligation of the MazF-truncated 16S rRNA terminus established the reversibility of one form of ribosomal RNA cleavage (*4*), while the RtcB2–PrfH system extended this principle to damage within the decoding center (*5*). RtcB has also been implicated in recovery of fragmented rRNA following antibiotic stress (*25*), and replacement of oxidized ribosomal proteins provides a distinct route to ribosome repair in yeast (*26*). Our structures reveal how an RNA repair ligase engages a damaged ribosomal subunit and capture covalent intermediates before and after guanylate transfer to its rRNA substrate. They therefore connect recognition of the damaged translation machinery to the chemistry of RNA-end activation.

The distinction between enzyme-linked GMP in states I/II and RNA-linked GMP in state III directly relates the structures to the established RtcB reaction pathway. Previous biochemical studies identified enzyme–GMP and RNA–GMP intermediates (*27*), whereas structural studies defined histidine guanylylation and used sulfate ions to infer substrate phosphate positions (*12, 15*). The present structures locate the A1493 phosphate acceptor beside H294-linked GMP and reveal the resulting RNA(3′)pp(5′)G intermediate within a largely preserved catalytic and metal-binding architecture. Unlike classical ATP-dependent ligases, which activate a 5′-phosphate for attack by a 3′-hydroxyl, RtcB activates a 3′-phosphate for attack by a 5′-hydroxyl (*6–8, 14, 16*). Nevertheless, activation alone is insufficient for strand joining. Step-resolved biochemical analysis implicated recognition of the hydroxyl-containing strand as a distinct catalytic requirement (*28*), and the PhoRtcB–DNA structure revealed contacts with that strand (*21*). Entry of the G1494-containing segment into cleft5 in state III provides a structural connection between activation of the ribosomal 3 ′-phosphate and accommodation of the opposite end before ligation.

Productive engagement involves both stabilization of the ribosomal scaffold and local rearrangement of the substrate. Reciprocal insertion of R52 into h44 and A1492 into *Ec*RtcB2 creates an interlocking interface near the break, while contacts with head-associated rRNA and uS12 extend recognition beyond the catalytic pocket. The broad variation in unbound head orientations contrasts with the closely similar head configurations in the three bound states, supporting stabilization of the head and overall h44 framework upon *Ec*RtcB2 binding. Within this framework, *Ec*RtcB2 reorients and the G1494-containing segment adopts a different configuration in the RNA–GMP state. Thus, scaffold stabilization is compatible with local changes associated with RNA activation. The structures associate these changes but do not establish whether enzyme reorientation promotes downstream-end entry or follows it.

The extended interface also offers an explanation for specialization within the RtcB family. Functional differences among bacterial paralogs were evident in earlier studies of *Myxococcus xanthus* RtcB proteins (*29*), and our previous work established distinct substrate preferences for *Ec*RtcB1 and *Ec*RtcB2 (*5*). The tRNA–h44 superposition reveals no apparent steric clashes with *Ec*RtcB2 in the modeled orientation, arguing against gross exclusion as a sufficient explanation for its inability to repair tRNA. Instead, productive recognition may depend on the combined geometry of peripheral contacts and reactive-end positioning. Differences at the head- and uS12-facing surfaces could impair d30S engagement by *Ec*RtcB1, whereas the longer loop near the 5′-hydroxyl-binding region of *Ec*RtcB1 and *Pho*RtcB identifies a potential contribution to accommodation of other substrates. These comparisons provide candidate determinants of specificity, without establishing their individual functional contributions or equating failure of ligation with failure of binding.

Access to the lesion imposes an additional constraint. The observed *Ec*RtcB2-binding mode overlaps with H69 and with ligands occupying the decoding region in the examined 70S configurations, while the d70S reconstruction retains structural barriers involving H69 and the head. Subunit separation therefore provides a route to expose the damaged decoding center after PrfH-mediated peptide release (*5, 22*). Repair detected in reactions initiated with mRNA- and P-site tRNA-containing 70S complexes (*23*) does not by itself establish the subunit-association state during ligation. Transient dissociation could expose a repairable d30S population, while alternative associated-ribosome conformations remain another possibility. Our structures define the incompatibility of the observed binding mode with the examined 70S states rather than excluding every possible route to repair within an associated particle.

Together, these findings support a structure-guided model of ribosome repair (**Fig. 4**). PrfH-mediated peptide release precedes a proposed subunit-separation step that permits access to d30S. *Ec*RtcB2 engagement stabilizes the head and h44 framework and positions the A1493 3′-phosphate beside H294-linked GMP, as captured in states I and II. Transfer of GMP to A1493 generates the activated RNA intermediate observed in state III, accompanied by enzyme reorientation and engagement of the G1494-containing segment in cleft5. Subsequent attack by the G1494 5 ′ -hydroxyl would restore the rRNA backbone and release GMP, followed by dissociation of *Ec*RtcB2 and return of the repaired subunit to translation. Although the final strand-joining geometry and product-release steps remain unresolved, the structures establish how recognition and chemical activation are coordinated during repair of a damaged ribosomal substrate.

**Fig. 4.**
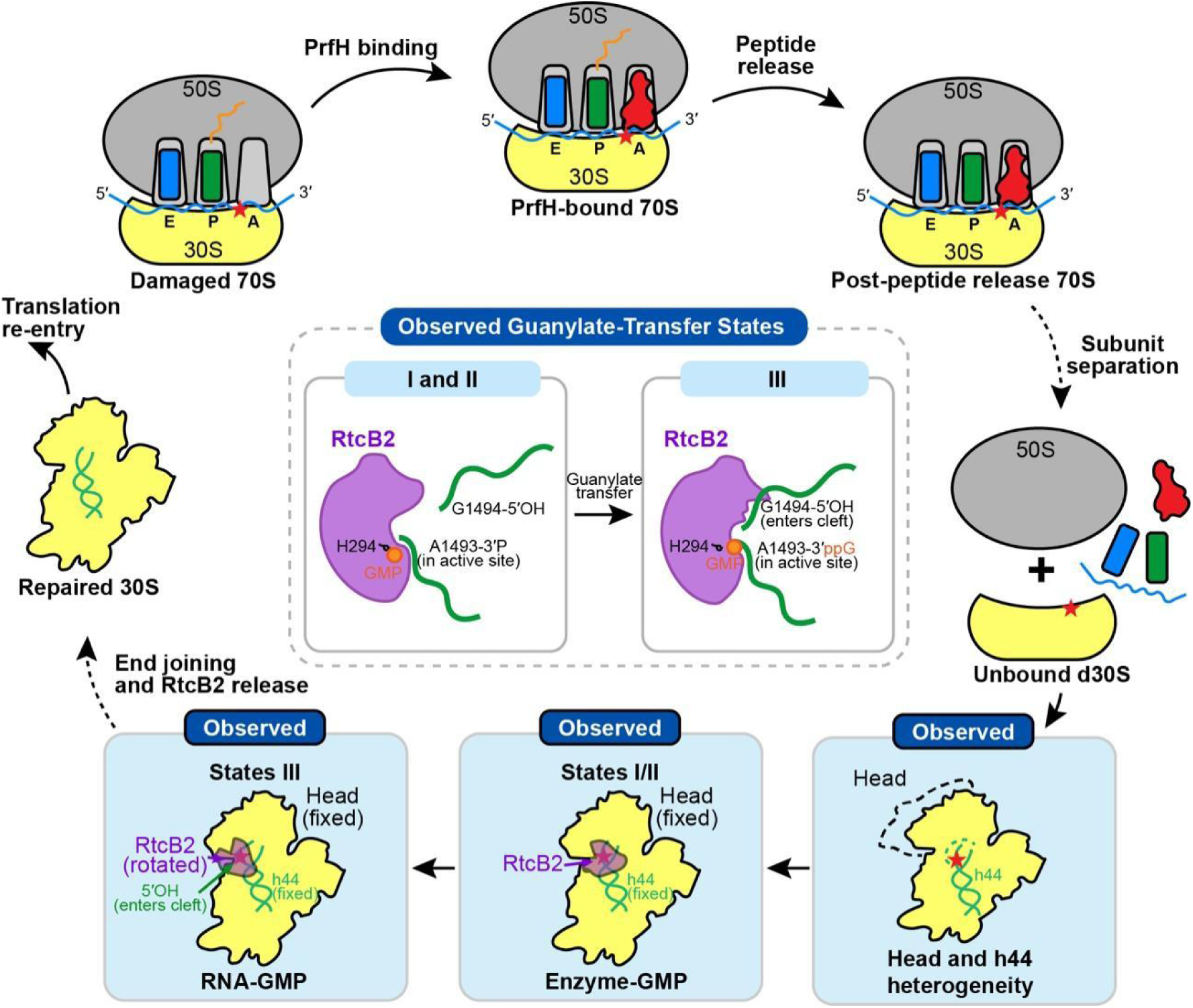
A Structure-guided model of RtcB2-mediated ribosome repair. PrfH recognizes the damaged 70S ribosome and promotes peptide release. Subsequent subunit separation exposes d30S for RtcB2 binding, which stabilizes the head and h44. States I and II contain H294-linked GMP with the A1493 3′-phosphate positioned in the active site. In state III, GMP is transferred to A1493, forming the RNA–GMP intermediate, while the G1494-containing 5′-hydroxyl segment enters an adjacent cleft. Proposed end joining and RtcB2 release restore the repaired 30S subunit for translation re-entry. Blue boxes highlight structures observed in this study; dashed arrows indicate proposed transitions. Red stars mark the rRNA cleavage site.

## MATERIALS AND METHODS

### Cloning, Expression, and Purification of Recombinant *Ec*RtcB2

The gene encoding *Ec*RtcB2 from *E. coli* ATCC 25922 was cloned into the pRSF-1 vector to generate an N-terminal His_6_–SUMO fusion. The construct was expressed in *E. coli* BL21(DE3), and protein expression was induced with 0.5 mM isopropyl β-D-1-thiogalactopyranoside (IPTG), followed by overnight incubation at 18°C. Cells were harvested and resuspended in lysis buffer containing 20 mM Tris-HCl (pH 8.0), 200 mM NaCl, and 5% glycerol, and disrupted using a French press. The lysate was clarified by centrifugation at 18,000 × g for 40 min at 4°C, followed by filtration through a 0.45-μm filter. The clarified lysate was applied to a HisTrap HP column (Cytiva, catalog no. 17524802), and bound proteins were eluted with an imidazole gradient. Fractions containing His_6_–SUMO–*Ec*RtcB2 were pooled and incubated with Ulp1 protease to remove the fusion tag. The cleavage mixture was subsequently passed through a second HisTrap HP column to separate untagged *Ec*RtcB2 from His-tagged species.

### Purification of Damaged *E. coli* 30S Subunits

Damaged *E. coli* 30S subunits were prepared from *E. coli* MG1655 carrying pCH450::cdiA-CT^ECL^ and pTrc99a::cdiI^ECL^. Cells were grown in Luria–Bertani (LB) medium to an OD_600_ of 0.4, and CdiA-CT^ECL^ expression was induced by adding arabinose to a final concentration of 0.2%. After 5 min of induction, cells were harvested and lysed in buffer containing 20 mM HEPES (pH 7.5), 6 mM MgCl_2_, 30 mM NH_4_Cl, and 4 mM β-mercaptoethanol, supplemented with 0.5 mM phenylmethylsulfonyl fluoride and 5.5 μg/mL DNase I. Damaged 70S ribosomes were purified following previously described procedures (*30*). Purified ribosomes were dissociated into 30S and 50S subunits by lowering the Mg^2+^ concentration and separated by sucrose gradient ultracentrifugation under low-Mg^2+^ conditions (*31*). Fractions corresponding to the damaged 30S subunits were collected and pooled for subsequent complex assembly.

### Cryo-EM Sample Preparation and Data Acquisition

The *Ec*RtcB2•d30S complex was reconstituted by incubating 0.2 μM damaged 30S subunits with 2 μM *Ec*RtcB2 at a 1:10 subunit-to-enzyme molar ratio for 10 min at 37°C. The assembly buffer contained 20 mM HEPES (pH 7.5), 5 mM MgCl_2_, 30 mM NH_4_Cl, 4 mM β-mercaptoethanol, 0.5 mM phenylmethylsulfonyl fluoride, and 0.5 mM MnCl_2_.

For vitrification, 2.5 μL of the reconstituted complex was applied to glow-discharged Quantifoil holey carbon grids coated with a thin continuous carbon film (catalog no. Q3100CR1.3). After a 30-s incubation, the grids were blotted for 3 s at a blot force setting of 5 using a Vitrobot Mark IV (FEI) maintained at 4°C and 100% relative humidity, and immediately plunge-frozen into liquid ethane.

Cryo-EM data were collected using a Titan Krios G3i transmission electron microscope (FEI; instrument identifier D3796) operated at an accelerating voltage of 300 kV and equipped with a Falcon III direct electron detector (FEI). Images were acquired at nominal defocus values ranging from −1.5 to −1.0 μm. A total of 8,345 movies were recorded, each comprising 30 frames with a total electron dose of 30 e^-^Å^-2^.

### Cryo-EM Image Processing, Classification, and Refinement

Beam-induced movie-frame motion was corrected using MotionCor2 (*32*). All subsequent image processing, including contrast transfer function (CTF) estimation, particle picking, two-dimensional (2D) and three-dimensional (3D) classification, and refinement, was performed in cryoSPARC v4 (*33*). The processing workflow is summarized in Fig. S1.

The three datasets were initially processed separately. Following CTF estimation, ribosomal particles were selected using the Blob Picker and extracted with a box size of 348 × 348 pixels. Three rounds of 2D classification were performed to remove non-ribosomal particles. The retained particles were subsequently separated into *Ec*RtcB2• d30S, unbound d30S, and d70S classes by 3D classification, using a reference generated from the 30S component of the intact *E. coli* 70S ribosome structure (PDB: 7K00) (*24*). Particles assigned to each class were then pooled across the three datasets, yielding 596,848 *Ec*RtcB2•d30S particles, 848,405 unbound d30S particles, and 173,309 d70S particles. Additional rounds of 2D and 3D classification were used to remove poorly resolved particles.

The *Ec*RtcB2•d30S particle pool was first subjected to global 3D classification to separate particles lacking *Ec*RtcB2. Classes containing *Ec*RtcB2 were further classified using a local mask encompassing the *Ec*RtcB2-binding region, which removed additional unbound d30S particles and resolved three related conformational states. These states exhibited modest differences in the positions and orientations of the 30S head and *Ec*RtcB2 and were refined to overall resolutions of 2.68, 2.65, and 2.76 Å, respectively.

All unbound d30S particles, including those recovered during classification of the *Ec*RtcB2•d30S particle pool, were combined and subjected to further 3D classification. This analysis resolved five conformational states with a broader range of head orientations than observed among the *Ec*RtcB2-bound states. Homogeneous refinement yielded reconstructions at overall resolutions of 2.65, 2.57, 2.63, 4.27 and 2.67 Å.

The d70S particle pool was subjected to an additional round of 2D classification to remove poorly resolved particles. An initial 70S reference was generated by ab initio reconstruction, and homogeneous refinement of the retained particles yielded a reconstruction of d70S without detectable bound *Ec*RtcB2 at an overall resolution of 2.59 Å.

Overall resolutions were determined using the gold-standard Fourier shell correlation (FSC) criterion of 0.143, calculated between two half-maps refined independently from separate halves of each particle dataset after application of a soft spherical mask (*34*). Local resolution was estimated in cryoSPARC (*33*).

### Model Building, Refinement and Validation

For model building and refinement, the initial models used were the cryo-EM structure of *Ec*PrfH in complex with the damaged ribosome (PDB: 7SA4) (*5*) and the Alphafold3-predicted model of *Ec*RtcB2 (*35*). The initial models were fitted into the density map using UCSF ChimeraX (*36*), and the models of *Ec*RtcB2, h44, 30S head were adjusted based on the map features using Coot (*37*). Further refinement was performed in Phenix (*38*). The final models were validated by the MolProbity (*39*), and the structures and maps were visualized using UCSF ChimeraX and PymoL.

## SUPPLEMENTARY MATERIALS

This PDF file includes:

Figs. S1 to S9

Tables 1 and S1

## Acknowledgments

F.Z. is an investigator of SUSTech Institute for Biological Electron Microscopy. The authors acknowledge the assistance of the cryoEM center at Southern University of Science and Technology for data collection. We thank the support of Center for Computational Science and Engineering at Southern University of Science and Technology for data analysis. This work was supported by grants from National Natural Science Foundation of China [82602342 for T.Y, 92169111 for F.Z., 32171200 for F.Z., 32100977 for F.Z.]; Natural Science Foundation of Fujian Province [2026J001437 for T.Y.]; Guangdong Innovative and Entrepreneurial Research Team Program [2021ZT09Y104 for F.Z.]; Shenzhen Science and Technology Program [JCYJ20220530115210023 for F.Z.]; Educational and Scientific Research Projects for Young and Middle-Aged Teachers of Fujian Province [JAT251041 for T.Y.]; Scientific Research Startup Foundation for High-Level Talents of Fujian Medical University [XRCZX2026004 for T.Y.].

## Author contributions

Writing—original draft: Y.T., Q.Z., T.Y., and F.Z. Conceptualization: Y.T., Q.Z., T.Y., and F.Z. Investigation: Y.T., Q.Z., Q.Y., Y.H., W.Z., J.H., T.Y., and F.Z. Writing-review and editing: Y.T., Q.Z., Q.Y., Y.H., W.Z., J.H., T.Y., and F.Z. Methodology: Y.T., Q.Z., T.Y., and F.Z. Resources: Y.T., Q.Z., T.Y., and F.Z. Fund acquisition: T.Y., and F.Z. Data curation: Y.T., Q.Z., T.Y., and F.Z. Validation: Q.Z., T.Y., and F.Z. Formal analysis: Q.Z., Y.H., T.Y., and F.Z. Supervision: T.Y., and F.Z. Project administration: Y.T., T.Y., and F.Z. Visualization: Q.Z., Y.H., and F.Z.

## Data and material availability

Electron microscopy maps have been deposited in the Electron Microscopy Data Bank under accession codes EMD-84910, EMD-84839, and EMD-84918 for the damaged 30S•*Ec*RtcB2-I, II, and III complex, respectively. EMD-84771 for the d70S reconstruction. EMD-84913, EMD-84914, EMD-84915, EMD-84916, and EMD-84917 for d30S states A to E, respectively. Coordinates have been deposited in the Protein Data Bank under accession codes 47WO, 47TQ, and 47WW for the damaged 30S•*Ec*RtcB2-I, II and III complex, respectively. All data and code needed to evaluate and reproduce the results in the paper are present in the paper and/or the Supplementary Materials.

## Supplementary Figures

**Fig. S1.**
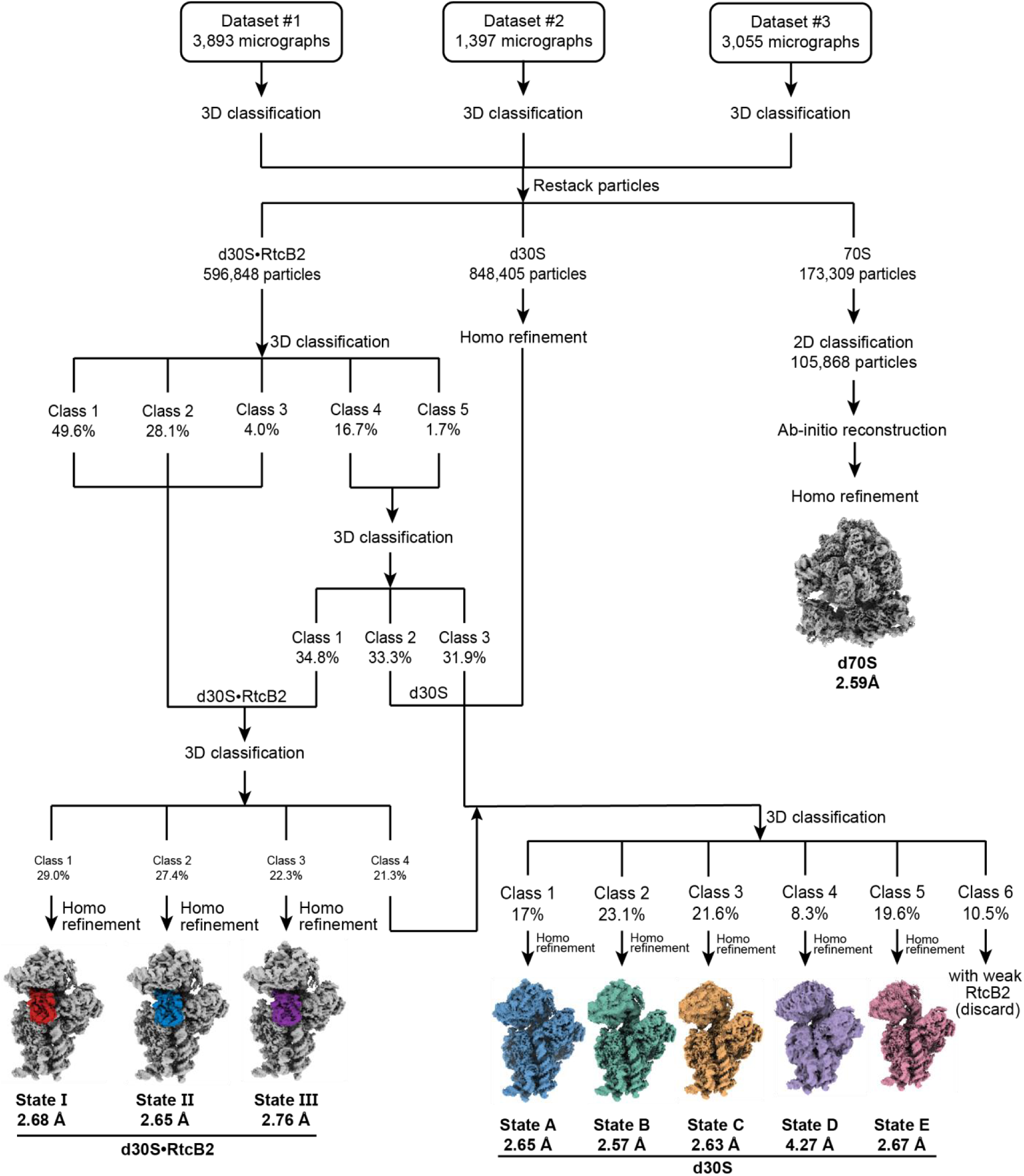
Cryo-EM classification of *Ec*RtcB2 in complex with the damaged *E.coli* 30S. Three datasets were merged in this study.

**Fig. S2.**
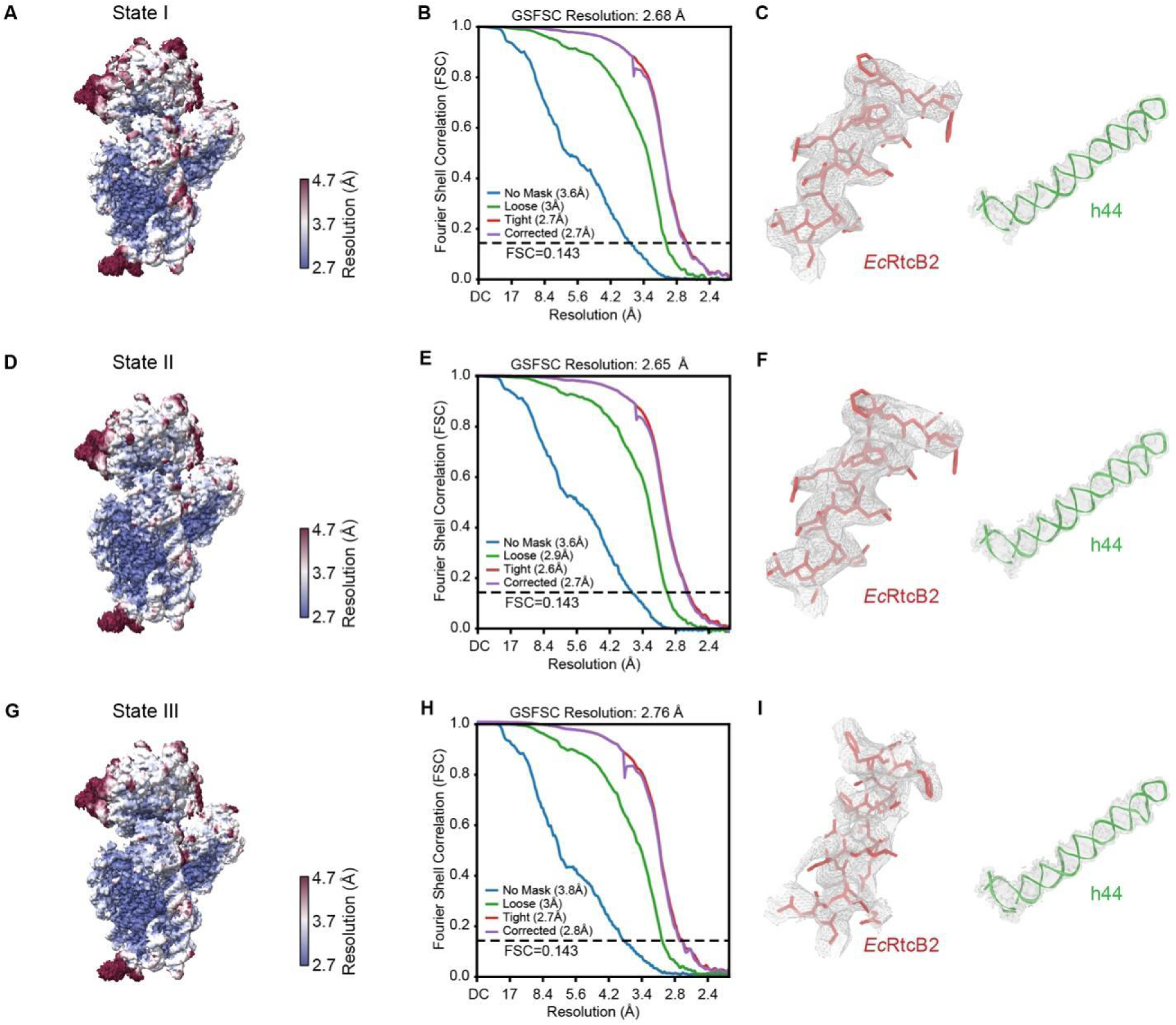
Map and model quality for the three states of d30S bound with *Ec*RtcB2. **A**, **D** and **G**, The maps coloured by local resolution in surface. **B**, **E** and **H**, Gold-standard FSC curves for the cryo-EM maps. **C**, **F** and I, Representative cryo-EM maps showing the refined structures of *Ec*RtcB2 (red) and h44 (green).

**Fig. S3.**
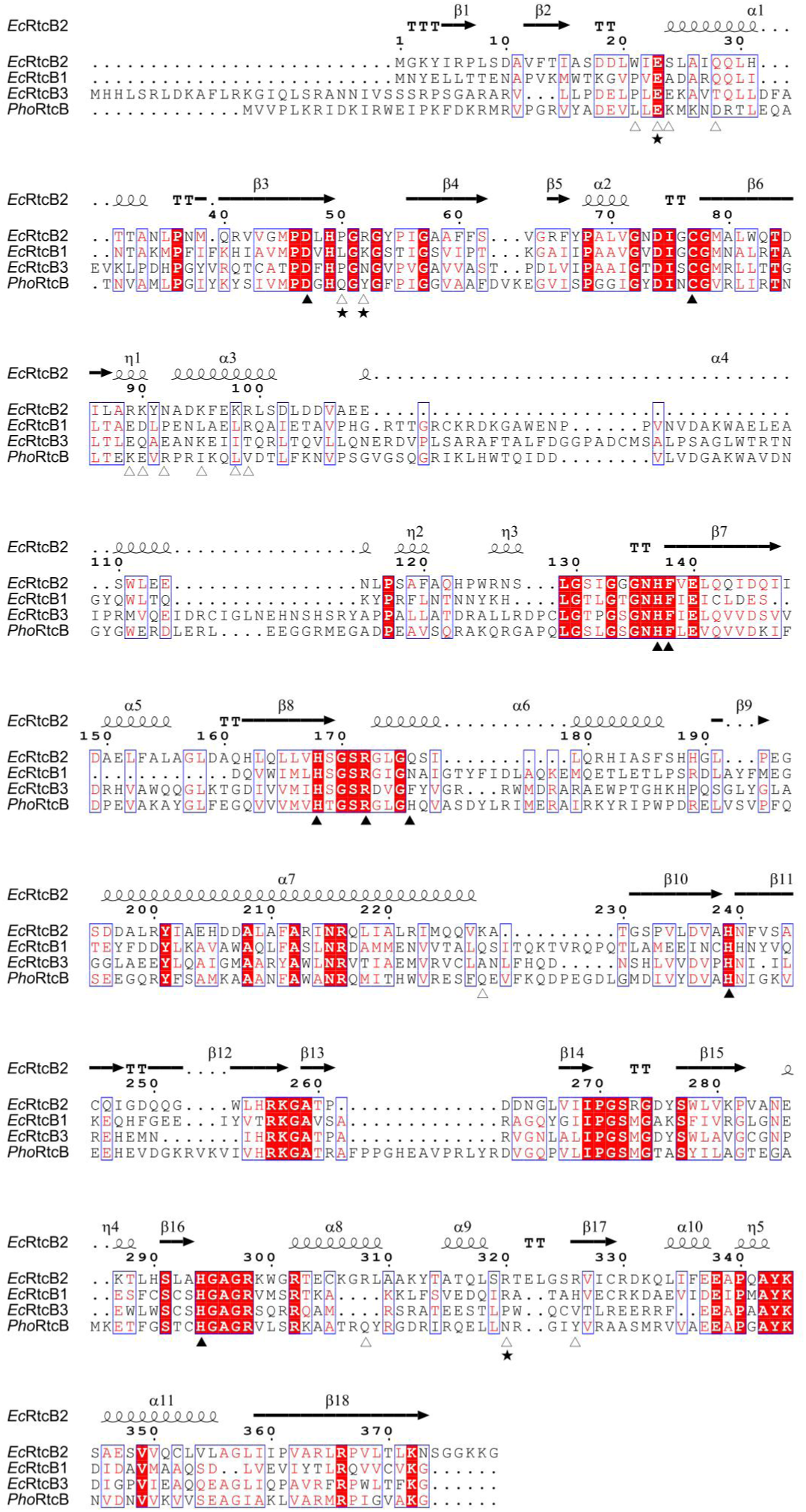
Sequence alignment of *Ec*RtcB2, *Ec*RtcB1, *Ec*RtcB3, and *Pho*RtcB. *Ec*RtcB2 secondary-structure elements and residue numbering are shown above the alignment. Filled triangles (▴) mark key active-site residues, and open triangles (△) indicate residues contacting 16S rRNA or uS12. Stars ( ★) additionally highlight P50, R52, R89, and R320, which pack against rRNA bases, and E23, which forms hydrogen-bonding contacts with A1492.

**Fig. S4.**
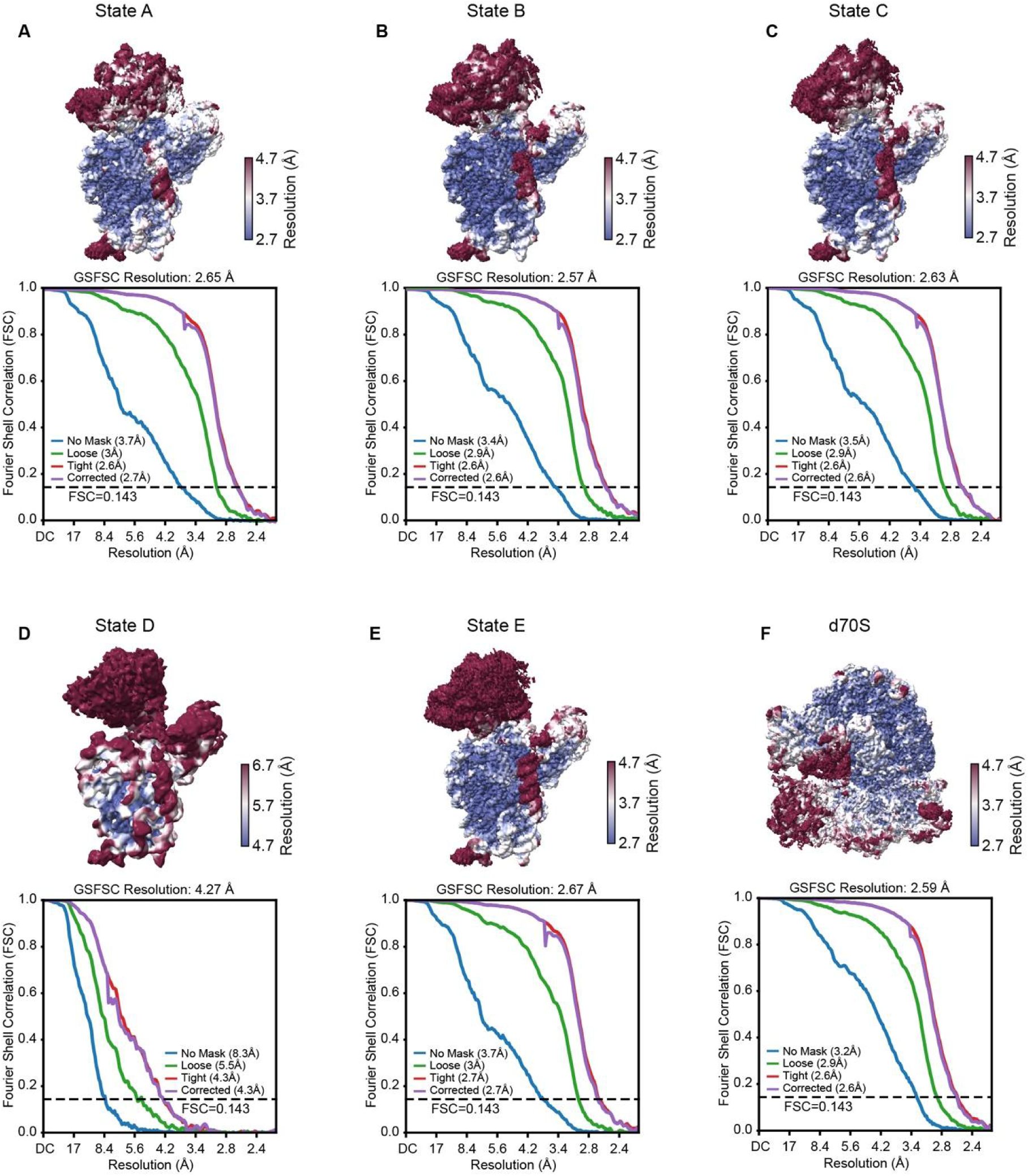
Map quality of the unbound d30S and d70S reconstructions. (A–E) Unbound d30S states A–**E**. (**F**) Damaged 70S ribosome. For each reconstruction, the upper panel shows a surface view of the cryo-EM map colored by local resolution, and the lower panel shows the gold-standard Fourier shell correlation (FSC) curves calculated without a mask (blue), with a loose mask (green), with a tight mask (red), and after correction for masking effects (purple).

**Fig. S5.**
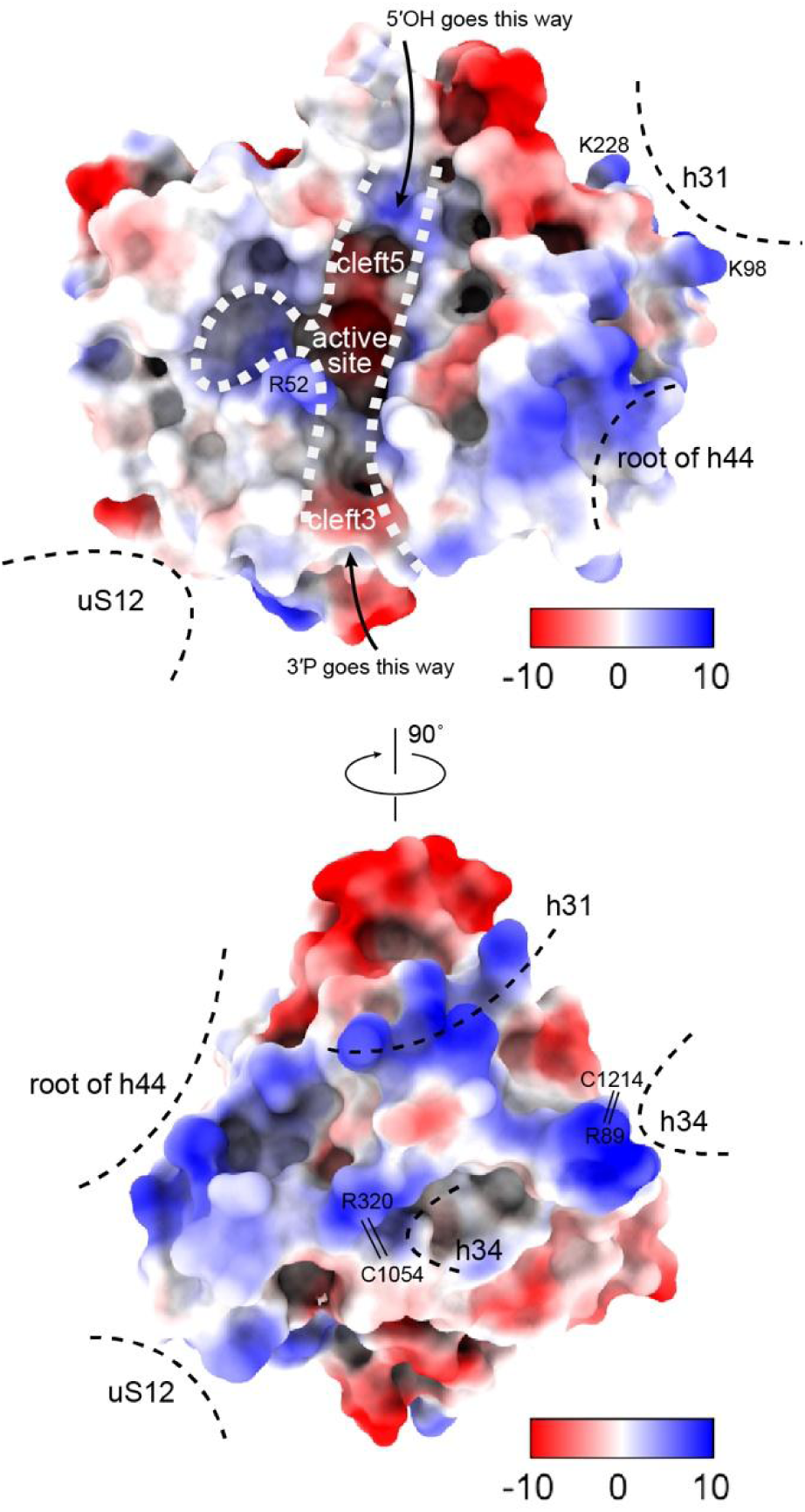
Electrostatic surface of *Ec*RtcB2. Two views related by a 90° rotation show positively charged patches at the rRNA-contacting surfaces. The clefts accommodating the 3′-phosphate and 5′-hydroxyl ends (cleft3 and cleft5) are partially electropositive. Dashed lines mark interaction regions, and arrows indicate RNA-end entry paths. The surface is colored by Coulombic electrostatic potential calculated in ChimeraX, ranging from −10 (red) to +10 (blue) kcal/(mol·e).

**Fig. S6.**
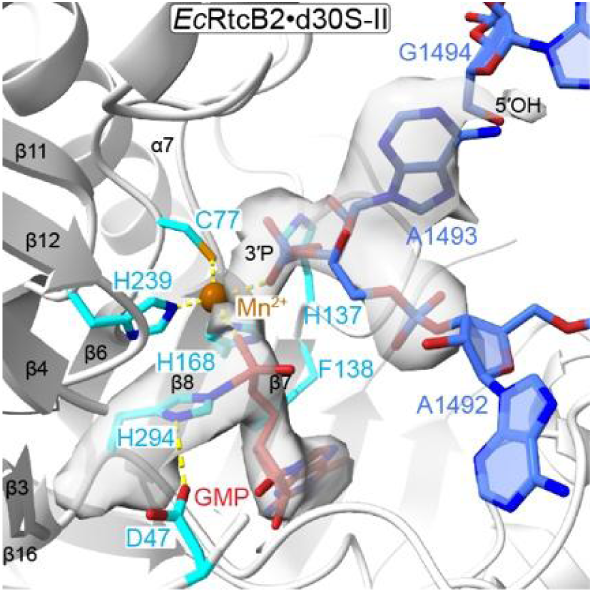
Active-site configuration of *Ec*RtcB2–d30S-II. Selected residues, rRNA nucleotides, and H294-linked GMP are shown as sticks with corresponding cryo-EM map features displayed as semitransparent surfaces. Mn^2+^ is shown as an orange sphere. The view corresponds to those in **Fig. 3A, B**.

**Fig. S7.**
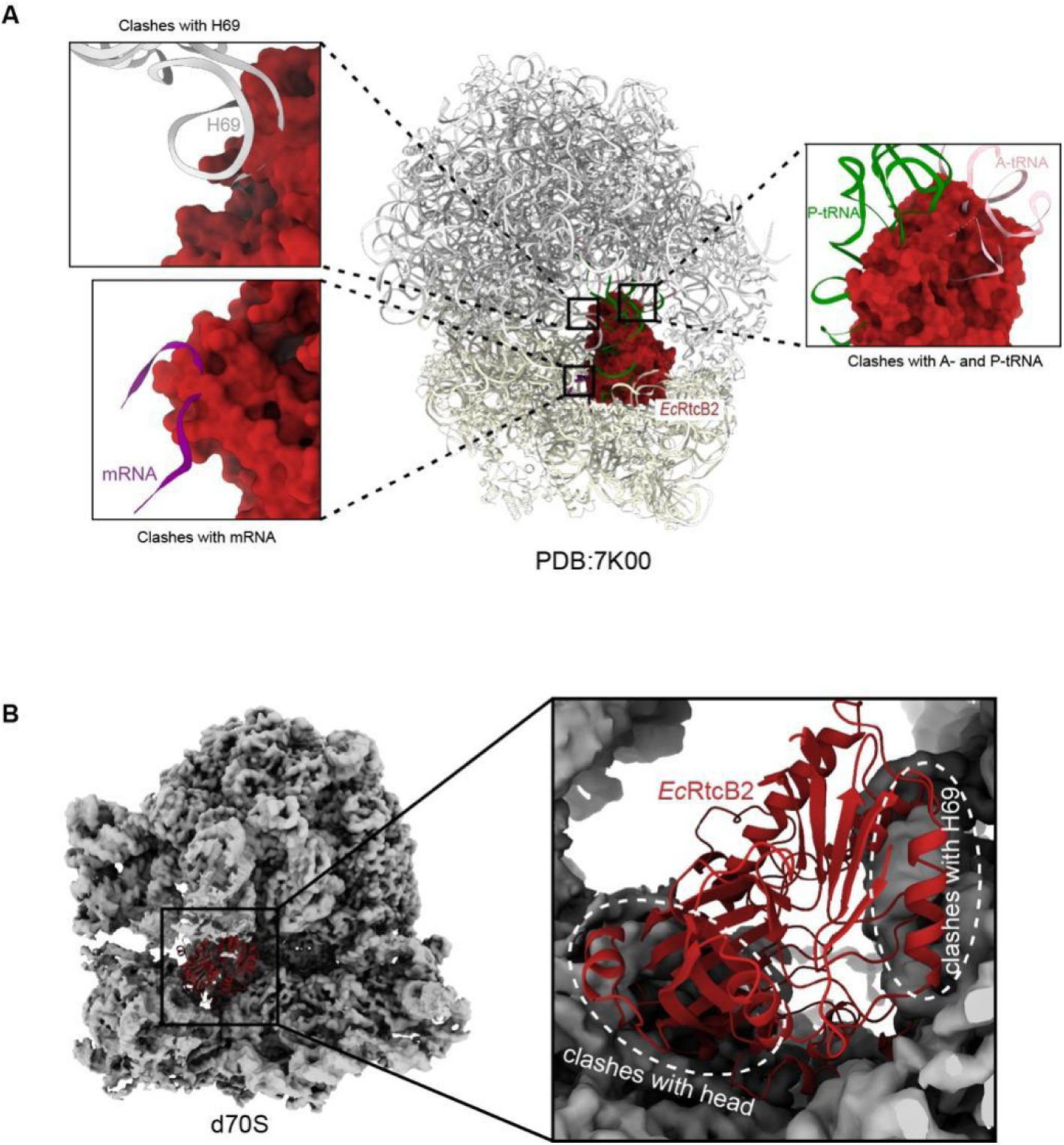
Structural constraints on *Ec*RtcB2 binding to assembled 70S ribosomes. (**A**) Superposition of the *Ec*RtcB2–d30S complex onto the *E. coli* 70S structure (PDB: 7K00) after alignment on the 30S body. Enlarged views show steric clashes of *Ec*RtcB2 (red surface) with H69, mRNA, and A- and P-site tRNAs. (**B**) Placement of *Ec*RtcB2 (red cartoon) into the d70S cryo-EM map by 30S-body alignment. The enlarged view highlights overlaps with H69 and the 30S head.

**Fig. S8.**
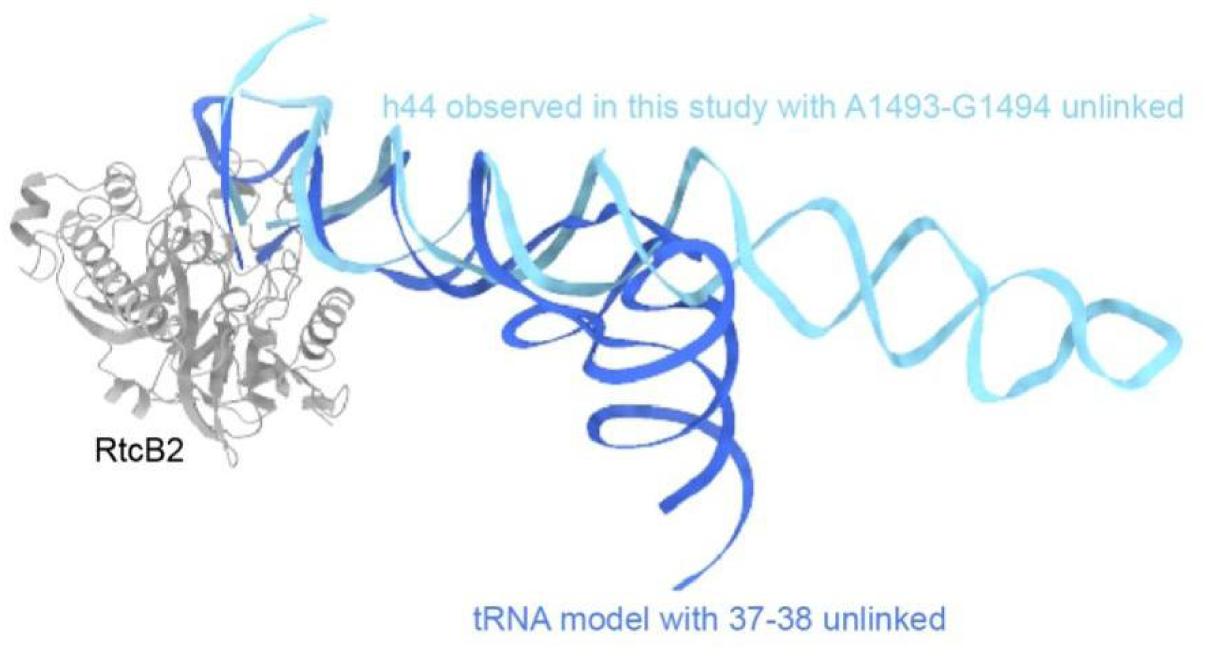
Structural superposition of a tRNA model and h44 bound to *Ec*RtcB2. A tRNA model with a break between nucleotides 37 and 38 (blue) is superimposed onto h44 cleaved between A1493 and G1494 (cyan). *Ec*RtcB2 is shown in gray. The modeled tRNA is accommodated without apparent steric clashes with *Ec*RtcB2. Model of tRNA was generated by alphaFold3.

**Fig. S9.**
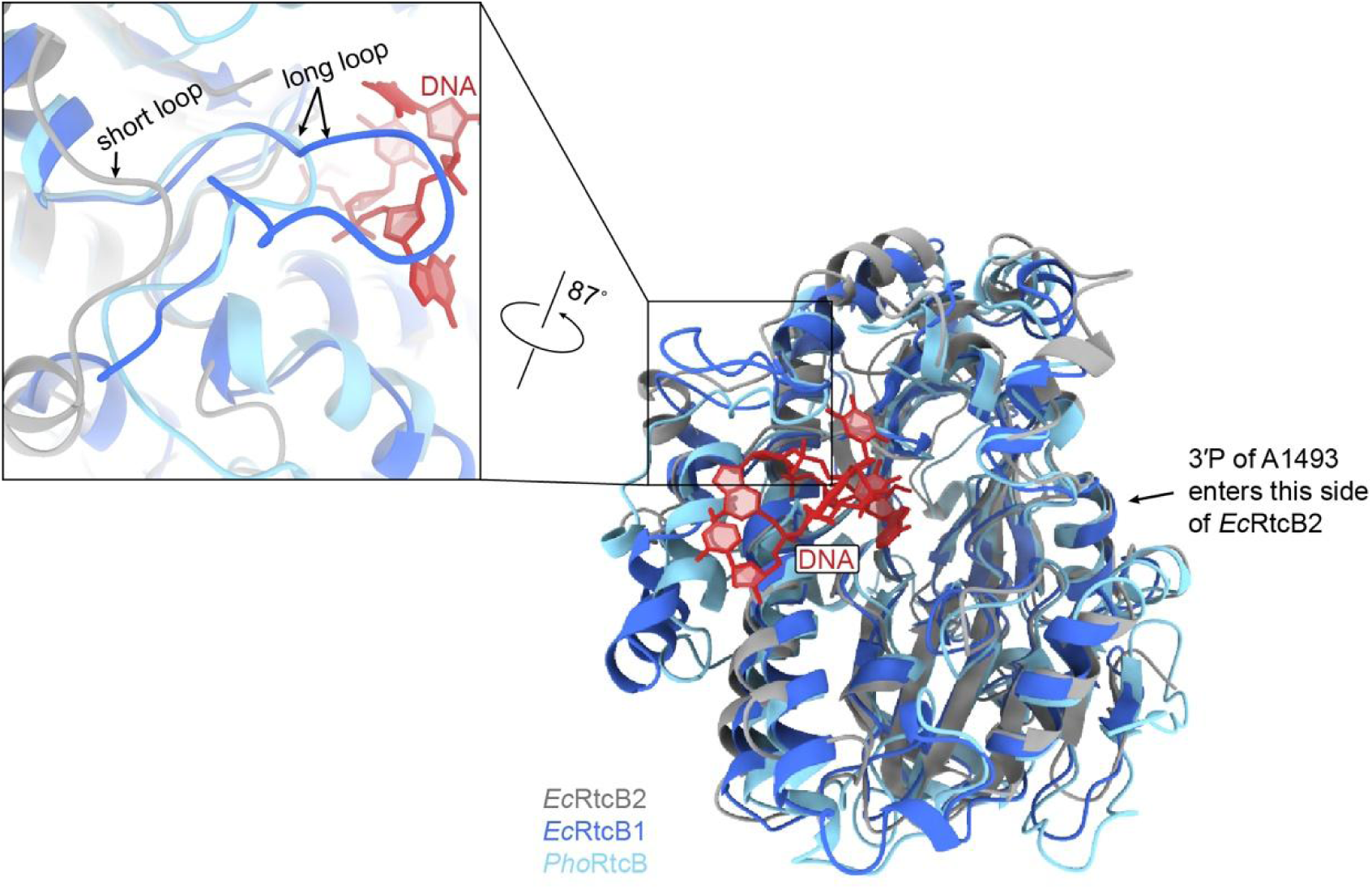
Structural comparison of *Ec*RtcB2, *Ec*RtcB1, and *Pho*RtcB. Superposition of *Ec*RtcB2 (gray), *Ec*RtcB1 (blue), and *Pho*RtcB (light blue), with the DNA oligonucleotide from the *Pho*RtcB complex (PDB: 7LFQ) shown in red. The proteins share a similar catalytic-pocket architecture but differ in an adjacent loop, which is shorter in *Ec*RtcB2. The inset highlights this region in a view rotated by 87°. The arrow indicates the side accommodating the A1493 3 ′ -phosphate in *Ec*RtcB2. Model of *Ec*RtcB1 was generated by alphaFold3.

**Table S1.** Data collection and model statistics for *Ec*RtcB2•d30S.

|  | ***Ec*RtcB2•d30S-I<br>(PDB:47WO)<br>(EMD-84910)** | ***Ec*RtcB2•d30S-II<br>(PDB: 47TQ)<br>(EMD-84839)** | ***Ec*RtcB2•d30S-III<br>(PDB:47WW)<br>(EMD-84918)** |
|---|---|---|---|
| **Data Collection** |  |  |  |
| Particles | 1,618,562 |  |  |
| Pixel size (Å) | 1.095 |  |  |
| Defocus range (µm) | -1.0 to -1.5 |  |  |
| Voltage (kV) | 300 |  |  |
| Electron dose (e<sup>-</sup>Å<sup>-2</sup>) | 30 |  |  |
| Particles for the reconstruction | 143,906 | 152,195 | 117,194 |
| **Model composition** |  |  |  |
| Non-hydrogen atoms | 54565 | 54565 | 54565 |
| Protein residues | 2782 | 2782 | 2782 |
| RNA bases | 1519 | 1519 | 1519 |
| Ligands (Mg<sup>2+</sup>/Mn<sup>2+</sup>) | 47/1 | 47/1 | 47/1 |
| **Refinement** |  |  |  |
| Resolution (Å) | 2.68 | 2.65 | 2.76 |
| Map sharpening B-factor (Å<sup>2</sup>) | 80.7 | 81.1 | 77.1 |
| Correlation coefficient against the maps | 0.91 | 0.91 | 0.91 |
| **Rms deviations** |  |  |  |
| Bond lengths (Å) | 0.009 | 0.009 | 0.011 |
| Bond angles (˚) | 0.788 | 0.759 | 0.864 |
| **Validation (proteins)** |  |  |  |
| Molprobity score | 1.71 | 1.66 | 1.78 |
| Clashscore, all atoms | 6.97 | 6.16 | 8.66 |
| Rotamer outlier (%) | 0.00 | 0.00 | 0.00 |
| **Ramachandran plot** |  |  |  |
| Favored (%) | 95.36 | 95.40 | 95.43 |
| Allowed (%) | 4.60 | 4.57 | 4.49 |
| Outlier (%) | 0.04 | 0.04 | 0.07 |
| **Peptide plane** |  |  |  |
| Cis proline/general (%) | 2.0/0.0 | 2.0/0.0 | 2.0/0.0 |
| Twisted proline/general (%) | 0.0/0.0 | 0.0/0.0 | 0.0/0.0 |

